# ‘Drifting’ *Buchnera* genomes track the microevolutionary trajectories of their aphid hosts

**DOI:** 10.1101/2023.11.17.567149

**Authors:** Joshua A. Thia, Dongwu Zhan, Katie Robinson, Paul A. Umina, Ary A. Hoffmann, Qiong Yang

## Abstract

Evolution of *Buchnera*–aphid host symbioses is often studied among species at macroevolutionary scales. Investigations within species offer a different perspective about how eco-evolutionary processes shape patterns of genetic variation at microevolutionary scales. Our study leverages new and publicly available whole-genome sequencing data to study *Buchnera*–aphid host evolution in *Myzus persicae*, the peach potato aphid, a globally invasive and polyphagous pest. Across 43 different asexual, clonally reproducing isofemale strains, we examined patterns of genomic covariation between *Buchnera* and their aphid host and considered the distribution of mutations in protein-coding regions of the *Buchnera* genome. We found *Buchnera* polymorphisms within aphid strains, suggesting the presence of genetically different *Buchnera* strains within the same clonal lineage. Genetic distance between pairs of *Buchnera* samples was positively correlated to genetic distance between their aphid hosts, indicating shared evolutionary histories. However, there was no segregation in the genetic variation of both *M. persicae* and *Buchnera* with plant host (Brassicaceae and non-tobacco Solanaceae) and no associations between genetic and geographic distance at global or regional spatial scales. Abundance patterns of non-synonymous mutations were similar to synonymous mutations in the *Buchnera* genome, and both mutation classes had similar site frequency spectra. We hypothesize that a predominance of neutral processes results in the *Buchnera* of *M. persicae* to simply ‘drift’ with the evolutionary trajectory of their aphid hosts. Our study presents a unique microevolutionary characterisation of *Buchnera*–aphid host genomic covariation across multiple aphid clones. This provides a new perspective on the eco-evolutionary processes generating and maintaining polymorphisms in a major pest aphid species and its obligate primary endosymbiont.

## INTRODUCTION

Endosymbiotic bacteria can play significant roles in the physiology of their host organisms. Among the most well studied endosymbionts is *Buchnera aphidicola* (herein, ‘*Buchnera*’), a maternally inherited proteobacteria found in aphids. *Buchnera* has a long evolutionary history with aphids, which has led to the two organisms being entwined in an obligate symbiosis. *Buchnera* have lost many core genes for their own regulation, but maintain genes involved in the biosynthesis of essential amino acid missing from the diet of their sap-sucking aphid hosts (Moran & Mira, 2001; Shigenobu, Watanabe, Hattori, Sakaki, & Ishikawa, 2000). Because of the importance of *Buchnera* to the fitness of its aphid host, it is often described as a primary endosymbiont. This contrasts with secondary endosymbionts, which are other non-obligate endosymbiotic bacteria that can have important, but variable, phenotypic effects on their aphid host (Guo et al., 2017). At broad macroevolutionary scales (among species), there is strong congruence between the phylogenies of aphid species and their *Buchnera*, reflecting unique coevolutionary histories of adaptation and (or) drift in the genomes of these organisms over evolutionary time (Chen, Wang, Chen, Jiang, & Qiao, 2017; Chong, Park, & Moran, 2019; Jousselin, Desdevises, & Coeur D’Acier, 2009; Nováková et al., 2013).

Coevolution between *Buchnera* and their aphid hosts can also be studied at the microevolutionary scale (within species), providing a population genetic perspective and an appreciation for processes that generate and maintain genetic polymorphisms in these symbioses over ecological time (Bennett & Moran, 2015; Wernegreen, 2015, 2017). To date, microevolutionary studies on the *Buchnera*−aphid host symbiosis have primarily focused on the model species, the pea aphid, *Acyrthosiphon pisum* (as reviewed in: Bennett and Moran 2015, Smith and Moran 2020). In *A. pisum*, specific genetic variants in the *Buchnera* genome have been linked to thermal tolerance of their aphid host (Dunbar, Wilson, Ferguson, & Moran, 2007; Perreau, Zhang, Maeda, Kirkpatrick, & Moran, 2021) and performance under diets depleted of essential amino acids, such as histidine (Chung, Parker, Blow, Brisson, & Douglas, 2020) and arginine (Vogel & Moran, 2011). Phylogenetic patterns among *Buchnera* genomes have been used to track plant host specialization of *A. pisum* across different species of Fabaceae (Peccoud, Simon, McLaughlin, & Moran, 2009). Other laboratory studies on *A. pisum* have helped develop insight into the (lack of) plasticity of *Buchnera* gene expression in response to nutrient variability (Moran, Plague, Sandström, & Wilcox, 2003) and possible interactions with gene expression patterns in their aphid host (Smith & Moran, 2020). Whilst *A. pisum* has provided a valuable system for understanding microevolutionary dynamics in the *Buchnera*–aphid host symbioses, we lack a broader perspective of how variation in *Buchnera* might affect the evolutionary ecology in other aphid species (including major pest species).

*Myzus persicae*, the peach potato aphid, is in many ways a prime species for studying coevolution between *Buchnera* and its aphid host on microevolutionary scales. *Myzus persicae* originated in Asia, but is now a major globally invasive agricultural pest (Li et al., 2015). This global distribution of *M. persicae* is interesting with respect to considering possible genomic changes that might be associated with post-invasion adaptation or drift. Clonal diversity in invasive ranges can be low, presumably through the joint action of post-invasion bottlenecks and strong selective sweeps from agricultural insecticides (de Little, Edwards, van Rooyen, Weeks, & Umina, 2017; Umina, Edwards, Carson, Van Rooyen, & Anderson, 2014). However, the extent to which related, but geographically separated, clones diverge from each other post-invasion has been poorly characterized, with previous work largely interested in quantifying beta diversity in clonal composition (e.g., Margaritopoulos, Kasprowicz, Malloch, & Fenton, 2009; Singh et al., 2021). *Myzus persicae* is also an incredibly polyphagous pest that is able to infest more than 40 different plant families (CABI Compendium, 2024); this strongly contrasts the model *A. pisum*, which is restricted to plants in the family Fabaceae (Peccoud et al., 2009). Unlike many other aphids species, *M. persicae* tends to lack secondary endosymbionts, especially in its invasive range (Yang et al., 2023). The rarity of these secondary endosymbionts in *M. persicae* would simplify the possible microevolutionary dynamics between the aphid host and *Buchnera*, as well as attempts to link genomic variation in their genomes to environmental variables.

We know very little about segregating patterns of *Buchnera* variation within and among different clonal lineages of *M. persicae*. Some prior work has suggested that the *Buchnera* of *M. persicae* exhibit a greater retention of genes important for nutritional symbioses (e.g., aspartate, methionine, and queosine production) relative to other aphid species, which may be linked to the polyphagous nature of their *M. persicae* host (Argandona, Kim, & Hansen, 2023; Douglas, 1988; Jiang et al., 2013). Prior global analysis of *M. persicae* clones revealed an association between the aphid host genome and different plant host families (Singh et al., 2021). In that study, the major axis of aphid host genetic structuring was related to the primary peach host and recent tobacco specialization, but genetic structuring with respect to other plant hosts was less clear, and structuring of genetic variation in *Buchnera* was not considered. Some additional work in tobacco specialized *M. persicae* clones suggests that adaptive genetic changes underlying this plant host shift have occurred in the aphid host rather than in the *Buchnera* genome (Jiang et al., 2013; Singh et al., 2020). A greater understanding of how *Buchnera* may (or may not) contribute to host plant specialization in *M. persicae* could be gained from examining a broader diversity of clones from other plant hosts.

Here, we investigate the *Buchnera*-aphid symbiosis in *M. persicae* on a microevolutionary scale. Our analyses focus on patterns of genomic covariation between the *Buchnera* and aphid host genome *within* clonal aphid strains and the structuring of this genomic covariation *among* clone strains across space and on different host plants. *Buchnera*–aphid host genomic covariation could arise from the near-neutral processes of mutation accumulation and genomic decay, so that lineage specific genomic changes in *Buchnera* simply track the lineage of their aphid host via drift (McCutcheon & Moran, 2012). Adaptive processes could also contribute to patterns of covariation, such as selection favoring coadaptations between the aphid host and endosymbiont that increase overall fitness (Gilbert et al., 2010), or antagonistic selection for adaptations and counter-adaptations in a ‘tug-of-war’ between these organisms (Bennett & Moran, 2015; Smith & Moran, 2020). Under coadaptation, genetic variants in both the endosymbiont and aphid host could rapidly spread through a population and predominate descendent lineages, or else drive codivergence among lineages under diversifying selection, such as in response to different host plants and climates.

Our study had two primary aims. Our first aim was to quantify genomic variation within, and covariation between, *Buchnera* and its aphid host with respect to geography and plant host. This allowed us to investigate how genomic (co)variation is segregating among different clonal strains, providing insight into post-invasion genomic changes and plant host specialization, respectively. Our second aim was to characterize the distribution of mutations in protein-coding regions across *Buchnera* from different clonal aphid strains. This allowed us to specifically test for the potential contributions of adaptive versus neutral processes in shaping the distribution of variation within the *Buchnera* genome. We compiled a globally distributed dataset of asexually reproducing clonal strains of *M. persicae* using publicly available sequencing data and new sequence data from Australia. We focused on clonal strains associated with two less well studies plant families in *M. persicae* host specialization: Brassicaceae (cruciferous vegetables) and Solanaceae (peppers). We excluded clonal strains from tobacco (also a member of Solanaceae) in light of evidence suggesting that specialization on this plant host is due to genomic changes in the aphid host (Jiang et al., 2013; Singh et al., 2020) and support for tobacco specialized clones representing a distinct subspecies (*M. p. nicotianae*) (Singh et al., 2021). Our study provides among the first major characterizations of genomic covariation between aphid clones and their primary endosymbiont at the microevolutionary scale.

## METHODOLOGY

### Sample collection and metadata curation

Our study used a global sample of *M. persicae* (*n* = 43), leveraging previously sequenced *M. persicae* clones from Singh et al. (2021) (*n* = 24) and newly sequenced clones collected in this study from Australia (*n* = 19). All clonal strains were derived from plant hosts belonging to the families Brassicaceae (*n* = 22) or Solanaceae (*n* = 21) and established as asexually reproducing isofemale strains in the laboratory. We treat strains as a unique clonal lineage consisting of females with (largely) the same genotype given that genomes were characterized within a few generations. No secondary endosymbionts were identified in these clonal strains, as determined bioinformatically for samples sourced from Singh et al. (2021), and by molecular screening for samples sequenced in this study (following 16S metabarcoding and qPCR protocols outlined in Yang et al., 2023).

For the 19 Australian *M. persicae* clonal strains, we took 5 individuals and extracted them as a pooled DNA sample using Qiagen’s DNeasy^®^ Blood and Tissue kit. Whole-genome library construction was performed by Novogene (Novogene, Hong Kong, China) with sequencing performed on an Illumina HiSeq paired-end 150 bp. Sequence data for the clonal strains from Singh et al. (2021) was also obtained from pooled individuals. Herein, clonal samples follow the naming convention ‘Country_Study_ID’, where ‘Country’ is a country-of-origin code, ‘Study’ is one of ‘Z’ for this present study or ‘S’ for Singh et al. (2021), and ‘ID’ is a unique identifier number. Sample information is recorded in supplementary Table S1, and the global distribution of samples is illustrated in Figure 1. We retrospectively obtained microsatellite genotypes for the majority of Australian clones sequenced in this study (*n* = 14) to align our genomic information against historical measures of clonal variation in Australia (de Little et al., 2017; Umina et al., 2014),

**Figure 1.**
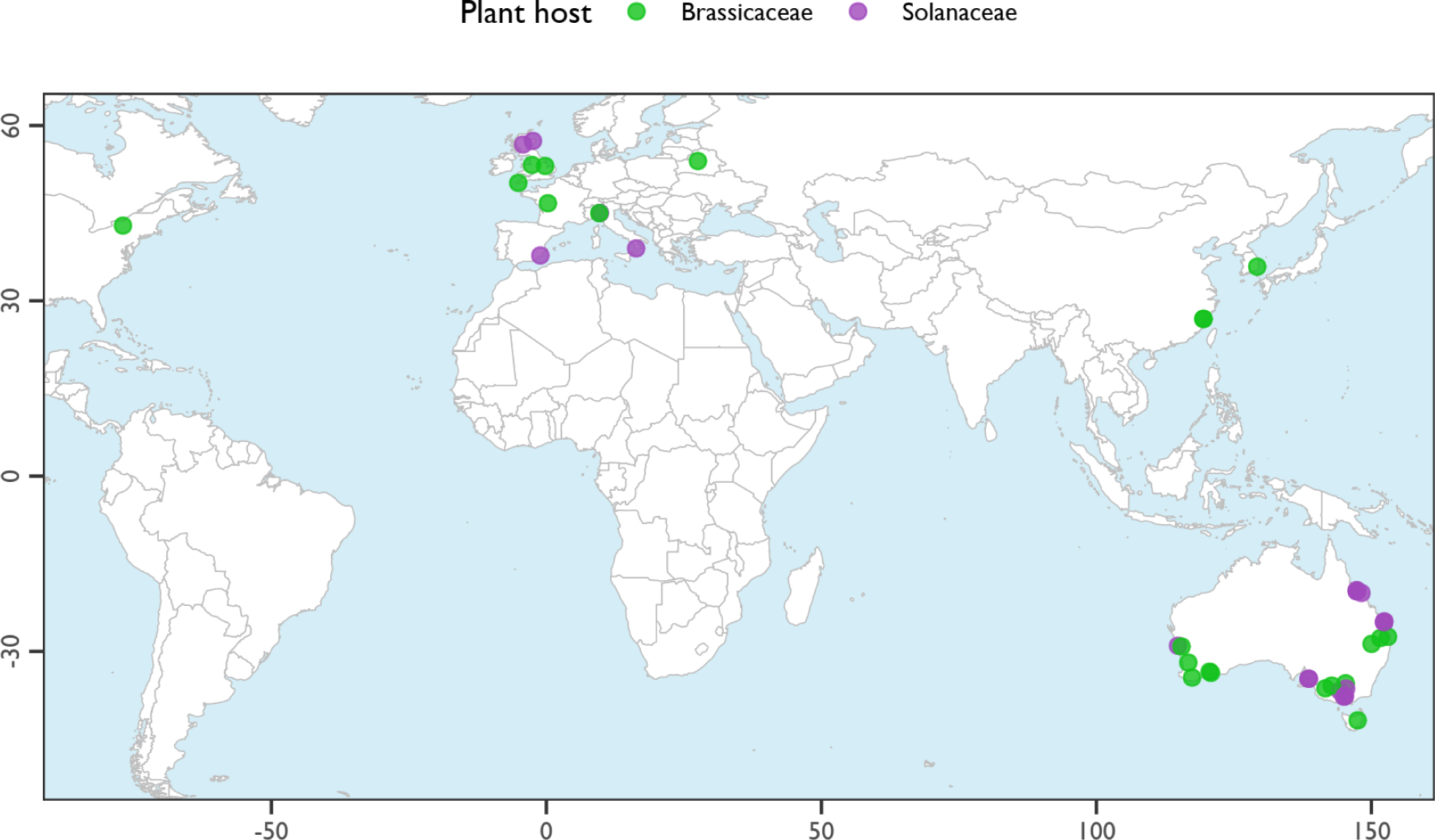
Global distribution of clonal *Myzus persicae* samples used in this study. Points are coloured by plant host family (see legend). See supplementary Table S1 for associated metadata.

### Read trimming, read mapping, and variant calling

Raw reads for our newly sequenced *M. persicae* and those from Singh et al. (2021) were trimmed and quality-filtered using *fastp* v0.12.4 (Chen et al., 2018). We then mapped trimmed reads against the G006 v3 nuclear reference genome for the aphid host (Aphid Base) and against the USDA strain *Buchnera* genome (GenBank Accession: CP002697), using *bowtie2* with the ‘very-sensitive’ end-to-end mode (Langmead et al., 2012). Alignments were sorted, filtered for a map quality >30, and deduplicated using *samtools* v1.16.1 (Danecek et al., 2021). The samples from Singh et al. (2021) had much greater sequencing depth than our low-coverage samples. We therefore down-sampled all Singh et al. (2021) reads to the maximum number of mapped reads observed for our samples to make the two datasets comparable: 15.6 million reads for *M. persicae* and 1.6 million reads for *Buchnera*. Down-sampling was performed with *samtools*. After down-sampling, we called variants in both the *M. persicae* host and *Buchnera* genomes using *freebayes* v1.3.2 (Garrison and Marth 2012). The raw variants was then separately filtered for analyses of (1) genetic diversity in the *M. persicae* host and *Buchnera* genomes, (2) genetic differentiation among *M. persicae* hosts and *Buchnera* genomes, and (3) mutational effects in the *Buchnera* genomes. We describe filtering for these analyses in the following subsections.

### Genetic diversity within clones

We quantified the genetic diversity in both the *M. persicae* host and *Buchnera* withon clones. We filtered each *Buchnera* and *M. persicae* clonal sample separately (*n* = 43 clones): (1) *Vcftools* v0.1.16 was used to subset variants to a minimum depth threshold (20 reads for *Buchnera* and 10 reads for *M. persicae* host genomes) and no missing data for each sample; (2) the number of SNP loci was extracted using custom code; (3) the total number of covered genomic sites meeting the minimum depth threshold was determined using the *samtools depth* function; (4) genetic diversity was calculated by dividing the number of observed SNPs in a clonal sample over the total covered genomic sites. In the context of *Buchnera*, this measure of genetic diversity represents the proportion of sites that are polymorphic across haploid genomes within an *M. persicae* clonal strain. In the context of the aphid host, this measure of genetic diversity represents the genomic heterozygosity of the diploid genome within a pooled sample from a *M. persicae* clonal strain. Because all pooled individuals were asexually produced, we treat the pool as representative (near approximation) of the diploid genotype.

We tested the hypothesis that genetic diversity differs between plant hosts. This allowed us to examine how genetic variation was structured across plant hosts. We fitted the model:

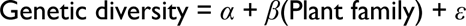

Here, the response is genetic diversity, *α* is the intercept, *ε* is the residual error, and *β* is regression coefficient for the effect of plant family (Brassicaceae or Solanaceae) as categorical variable. We used the *lm* function to fit model and the *anova* function to determine significance. Genetic diversity was transformed using a log10 transformation and addition of a small constant (1e−6) to reduce skew and non-normality, and to allow transformation of 0 values.

### Genetic differentiation between clones

We tested several hypotheses on the structuring of genetic variation between clones with respect to genetic differences across space and plant hosts. We filtered *Buchnera* and *M. persicae* host SNPs across all clonal samples following a series of filtering steps for all *n* = 43 clones: (1) *Vcftools* was used to reduce the variants to biallelic loci with a minimum quality of 20; (2) we manually reduced the data to SNPs using string searching on the VCF file; and (3) we used *vcftools* to filter out SNPs with minimum depth <20 reads across samples and any SNPs with missing data. SNPs were imported into R using the *genomalicious* R package (Thia and Riginos 2019) with the *vcf2DT* function. We performed a series of iterative filtering steps to more carefully curate genetic variants for downstream analysis, which involved: (1) setting a minimum read depth threshold, (2) filtering by a mean minor allele frequency of 0.05 across all samples, and (3) filtering to retain SNPs at a minimal distance threshold to reduce non-independence of sites. For *M. persicae* hosts, we set a minimum depth of 5 reads and a minimum SNP distance of 10,000 bp. For *Buchnera*, we set a minimum depth of 20 reads and a minimum SNP distance of 500 bp. This left 541 SNPs for the *M. persicae* host and 285 SNPs for *Buchnera* to estimate differentiation across clonal strains.

For *Buchnera*, we calculated the allele frequencies of the alternate allele by dividing the number of alternate allele reads by the total read count. However, SNPs with singleton reads for either the alternate or reference allele were set to a frequency of 0 or1, respectively, because singleton reads could be sequencing errors. Characterisation of allele frequencies treats the *Buchnera* as coexisting strains that coexist within a single clonal lineage of their host. Such a method has been used elsewhere to study genetic variation of coral endosymbionts (Matias et al., 2022), and other studies have used deep amplicon sequencing to study different strains of *Buchnera* in *A. pisum* (Perreau et al., 2021).

For *M. persicae*, we calculated genotype probabilities to help account for uncertainty in genotypes attributed to low-sequencing coverage (Clark et al. 2019). We performed Bayesian estimation of the genotype probabilities for each sample using the R package *polyRAD* (Clark et al. 2019). *PolyRAD* was designed in the context of polyploids but flexibly extends to diploids. Over-dispersion was examined using the *TestOverdispersion* function, with the optimal value for the over-dispersion parameter estimated as 26. The *IteratePopStructLD* function incorporated this over-dispersion into estimates with information on linkage, inferences of population structure, and expectations under Hardy-Weinberg equilibrium. Although the assumption of Hardy-Weinberg does not hold true for asexually reproducing clonal aphids, this approach allowed us to objectively infer genotype probabilities from low-coverage data as a proxy for diploid genotypes in clonal samples of the aphid host (Figure S1).

Next, we estimated pairwise differentiation across all clonal sample pairs. We used the Δ_D_ statistic as our measure of genetic differentiation, which measures beta diversity from relative abundance data based on Shannon’s entropy (Hill number of order *q* = 1), weighting alleles by their relative frequency. This deviates from genetic differentiation measured as *F*_ST_ (Hill number of order *q* = 2) which is influenced by heterozygosity rate (Gaggiotti et al., 2018). Additionally, Δ_D_ does not have the same underlying population genetic assumptions as *F*_ST_, and is arguably more appropriate for asexually reproducing species and for measuring allele frequency variation in *Buchnera* samples with haploid genomes. Δ_D_ was calculated using R’s IDIP function from the *HierDpart* package, available through GitHub (Gaggiotti et al., 2018). We produced a set of pairwise Δ_D_ values for *Buchnera* and another set for their *M. persicae* hosts.

Our first hypothesis tested for associations between genetic and geographic distance between pairs of clones. This hypothesis specifically tested whether sampled clones are increasingly genetically differentiated with increasing geographic distance. Geographically distant clones might be expected to diverge due to drift and (or) different local environments. Geographic distances between clonal samples were obtained using the *geosphere* package (Hijmans, 2022). We then correlated pairwise geographic distance against pairwise genetic differentiation, measured as Δ_D_, in the *Buchnera* and *M. persicae* host genomes.

Second, we tested whether patterns of differentiation between pairs of *M. persicae* genomes and pairs of their *Buchnera* genomes covaried. Such covariation could arise from both neutral and adaptive processes producing joint differentiation among *M. persicae* hosts and their *Buchnera*. We correlated pairwise Δ_D_ values between *Buchnera* and *M. persicae* genomes. To visualize coevolutionary patterns, we constructed neighbor joining trees for *Buchnera* and *M. persicae* from Δ_D_ distance matrices using the *nj* function from R’s *ape* package (Paradis, Claude, & Strimmer, 2004). Bootstrapping of node support was performed using *ape*’s *boot.phylo* function in conjunction with custom code to regenerate bootstrapped Δ_D_ distance matrices. Bootstrapping the Δ_D_ distance matrices required randomly redrawing SNP loci with replacement and then recalculating pairwise Δ_D_ values. Trees were constructed using R’s *ggtree* (Yu, Smith, Zhu, Guan, & Lam, 2017) and *treeio* (Wang et al., 2020) packages.

Third, we tested for broad evolutionary signals of plant host specialization. If *Buchnera* or *M. persicae* on the same plant host are genetically similar, pairwise genetic differentiation should be lower when comparing clones within a plant host relative to between plant hosts. We would also expect clustering of clonal strains by host plant. To test this, we first reduced our dataset to remove clonal lineages that were highly similar and found on the same plant. Our neighbour joining tree (see Results) showed that some samples formed clades that likely reflect groups of very closely related clones connected by short branch lengths. This included clonal samples from Australia, China, and the UK with pairwise Δ_D_ values ≤ 0.037. For those groups of closely related clones, we took the most derived sample within each clade to represent each plant host. This effectively acted as a phylogenetic correction to reduce pseudo-replication that might otherwise overinflate estimates of similarity within plant hosts. We were left with *n* = 25 clonal strains at the global scale, *n* = 11 clonal strains within Europe, and *n* = 12 clonal strains within Australia.

We tested our plant host specialization hypothesis using an analysis of residuals. We first fitted the models:

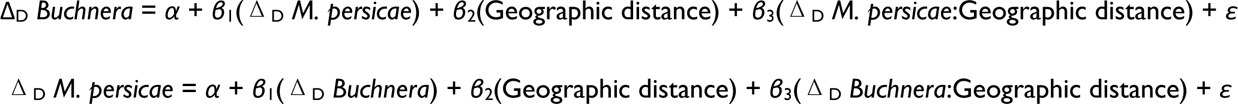

Here, the response is the pairwise genetic differentiation in a focal organism between clonal strains, *α* is the intercept, *ε* is the residual error, and *β*_1_, *β*_2_, *β*_3_ represent the partial regression coefficients of the partner organism’s pairwise genetic distance, pairwise geographic distance, and their interaction, respectively. All predictors were continuous. We then took the residuals and fitted two more models:

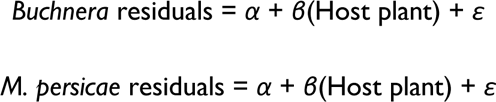

Here, the response is residual variation in pairwise genetic differentiation (unaccounted for by the partner’s genetic differentiation or geographic distance), *α* is the intercept, *ε* is the residual error, and *β* is the regression coefficient for host plant family, a categorical predictor with three levels (within Brassicaceae or Solanaceae, or between ‘different’ plant families). If there was a genome-wide signal of plant host specialization, we would expect residuals within Brassicaceae and Solanaceae samples to be smaller than those from different host plant samples. Models were fitted with R’s *lm* function with significance for each model tested using the *anova* function.

We tested each of our three hypotheses of genetic differentiation (genetic versus geographic distance, covariation between *Buchnera* and *M. periscae* hosts, and plant host specialization) using three different partitions of our dataset: global, Europe, and Australia. The global dataset provided a broad perspective but is complicated by potentially varying ecological factors among geographic regions and different evolutionary histories of clonal aphid lineages. We also had sufficient replication of samples from Europe and Australia, to perform these analyses separately in each of these regions to assess whether regional patterns differed from those observed at the global scale.

### Mutations in *Buchnera* protein-coding regions within and among clones

We examined protein-coding mutations and their distributions among *Buchnera* from different clonal samples to assess the possible relative contributions of adaptive versus neutral processes shaping genetic variation across this haploid genome. We filtered the raw variants with *vcftools* to retain any variant that was biallelic, had a minimum quality of 20, a minimum depth of 20, and no missing data. This dataset contained all variant types to capture the full range of segregating mutations among *Buchnera* genomes. Putative protein-coding effects were annotated with the program *snpEff* (Cingolani et al., 2012) and imported back into R. Variants that were annotated as ‘Missense’ (amino acid changes), ‘Stop’ (premature stop codon), or ‘Frameshift’ (altering reading frame) mutations were designated as ‘Non-synonymous’ mutations and compared with those identified as ‘Synonymous’ mutations. It is important to note that the number and class of mutations depends on the reference genome. We did not use an ancestral reference sequence, so mutations cannot be considered as being derived *per se.* Instead, we consider counts of alternate alleles. This approach provides insight into the relative importance of non-synonymous vs synonymous differentiation in protein-coding regions among *Buchnera* genomes from the same aphid host species.

We created site-frequency spectra (SFS) for non-synonymous and synonymous variants to test for similar frequency distributions between these two mutation classes. Downward projection was used to place counts of alternate alleles into bins from ‘1’ to ‘20’ by rounding the product of *q* × 20, where *q* is the alternate allele frequency. Prior to downward projection, any allele with a count of 1 read was set to a frequency of 0 to reduce the effect of singleton reads. Pearson’s correlation between allele count bins for non-synonymous versus synonymous mutations was calculated using the *ppcor* package (Kim, 2015).

We tested for enrichment (proportional over-representation) and diminishment (proportional under-representation) of non-synonymous variants in genes contributing to different functional pathways. We had four hypotheses. First, our ‘core cellular’ pathway hypothesis posited that genes involved in core cellular pathways should have fewer non-synonymous mutations that might reduce functionality. Decay of core cellular pathway genes would be detrimental to *Buchnera*. Second, our ‘core mutualism’ pathway hypothesis posited that genes involved in vital pathways to the *Buchnera*–aphid host symbiosis (e.g., amino acid biosynthesis) should have fewer non-synonymous mutations that might reduce functionality. Decay of core mutualism genes would be detrimental to the *M. persica* host. Third, our ‘ecologically relevant’ pathway hypothesis posited that genes involved in environmental adaptation should be enriched for non-synonymous mutations. Polymorphisms at ecologically relevant genes would benefit *Buchnera* by enhancing *M. persicae* host fitness under different ecological conditions. Fourth, our ‘drift’ hypothesis posited that neutral evolution predominates, such that non-synonymous and synonymous mutations occur at similar abundance across the *Buchnera* genome. In that case, no functional pathways would be expected to exhibit enrichment or diminishment of non-synonymous mutations.

Variants were assigned to KEGG functional pathways using the R packages *KEGGREST* (Tenenbaum, 2023) and *pathfindr* (Ulgen, Ozisik, & Sezerman, 2019). Contingency tables (2×2) were constructed for each focal *Buchnera* KEGG pathway: focal and non-focal genes (those genes in vs. not in the focal KEGG pathway, respectively) in rows, and non-synonymous and synonymous mutations in columns. Fisher’s exact test was used to test whether non-synonymous mutations were enriched or diminished in the focal functional pathway, using R’s *fisher.test* function with a two-sided test. Note, this analysis was performed using variants across all samples, and was not performed separately for each sample, to ensure a sufficient sample size and to obtain average effects across clonal strains.

## RESULTS

### Bioinformatic summary

The mean number of mapped reads to the haploid *Buchnera* genome was 616,927 with a range of 314,589 to 1,614,489 for our sequenced samples, and 3,089,599 with a range of 1,681,159 to 11,992,629 for the Singh et al. (2021) samples. The number of mapped reads to the aphid host diploid genome was 12.8 million with a range of 8.9 million to 15.6 million for our newly sequenced samples, and 79.1 million with a range of 38.1 million to 219.4 million for the Singh et al. (2021) samples.

### Genetic diversity within clones

For *Buchnera*, the mean proportion of polymorphic sites was 4.6e–5 with a range of 0 to 4.4e–4. For *M. persicae*, the genomic heterozygosity had a mean of 0.001 with a range of 0.0008 to 0.002 (Figure 2). There were no significant differences in genetic diversity between plant hosts in either *Buchnera or M. persicae* (Figure 2). *Buchnera* clearly exhibited variance in genomic the proportion of polymorphic sites across clones, which spanned several orders of magnitude. In contrast, the genomic heterozygosity in *M. persicae* had a smaller variance across clones. Importantly, that we observed polymorphism across haploid *Buchnera* genomes within clonal lineages suggests that there are different *Buchnera* strains segregating in aphid hosts from the same strain.

**Figure 2.**
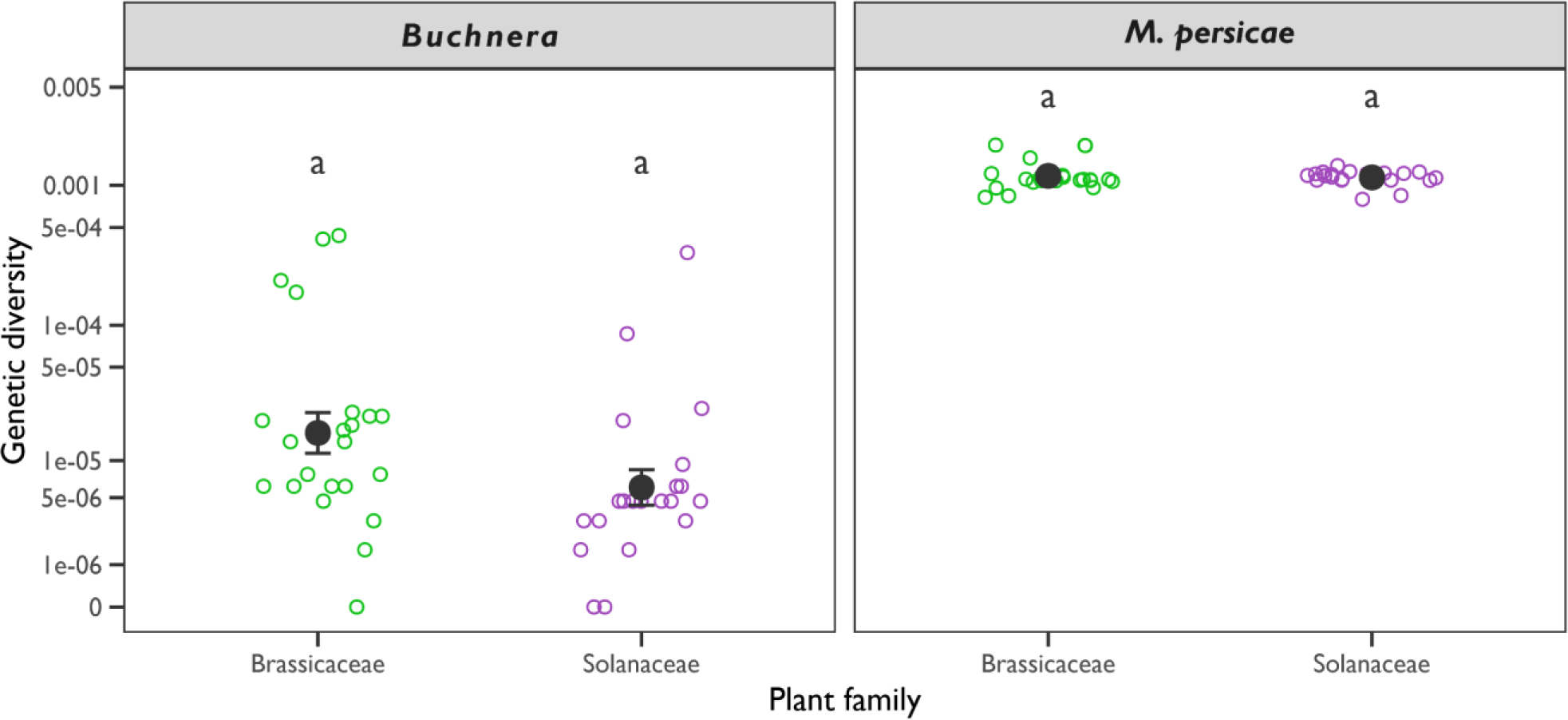
Patterns of genetic diversity in *Buchnera* and their *Myzus persicae* hosts. Genetic diversity within clones (*y*-axis) was measured as a function of host plant (*x*-axis). For *Buchnera*, genetic diversity is the proportion of sites that are polymorphic across haploid genomes within a clonal strain. For the *M. persicae* host, genetic diversity is the genomic heterozygosity of the diploid genome within a clonal strain. Open points represent clonal strains, large closed circles represent the mean with standard error bars. Panels separate estimates for *Buchnera* (left) and *M. persicae* hosts (right). Letters delineate significance group for the hypothesis that genetic diversity differs between plant hosts.

### Genetic differentiation between clones

Our global SNP set comprised 285 loci for *Buchnera* and 534 loci for *M. persicae*. Pairwise Δ_D_ values among *Buchnera* samples had a mean of 0.50, with a range of 0.0003 to 0.95. Pairwise Δ_D_ values among *M. persicae* samples had a mean of 0.27, with a range of 0.02 to 0.46. We found no evidence supporting associations between genetic and geographic distance in either *Buchnera* or *M. persicae* at the global scale, within Europe, or within Australia (Figures 3a, b, d, e, g, and h). Indeed, these patterns showed no obvious relationship; clonal strains separated by shorter geographic distances could be highly differentiated, and conversely, clonal strains separated by larger geographic distances could be genetically similar.

**Figure 3.**
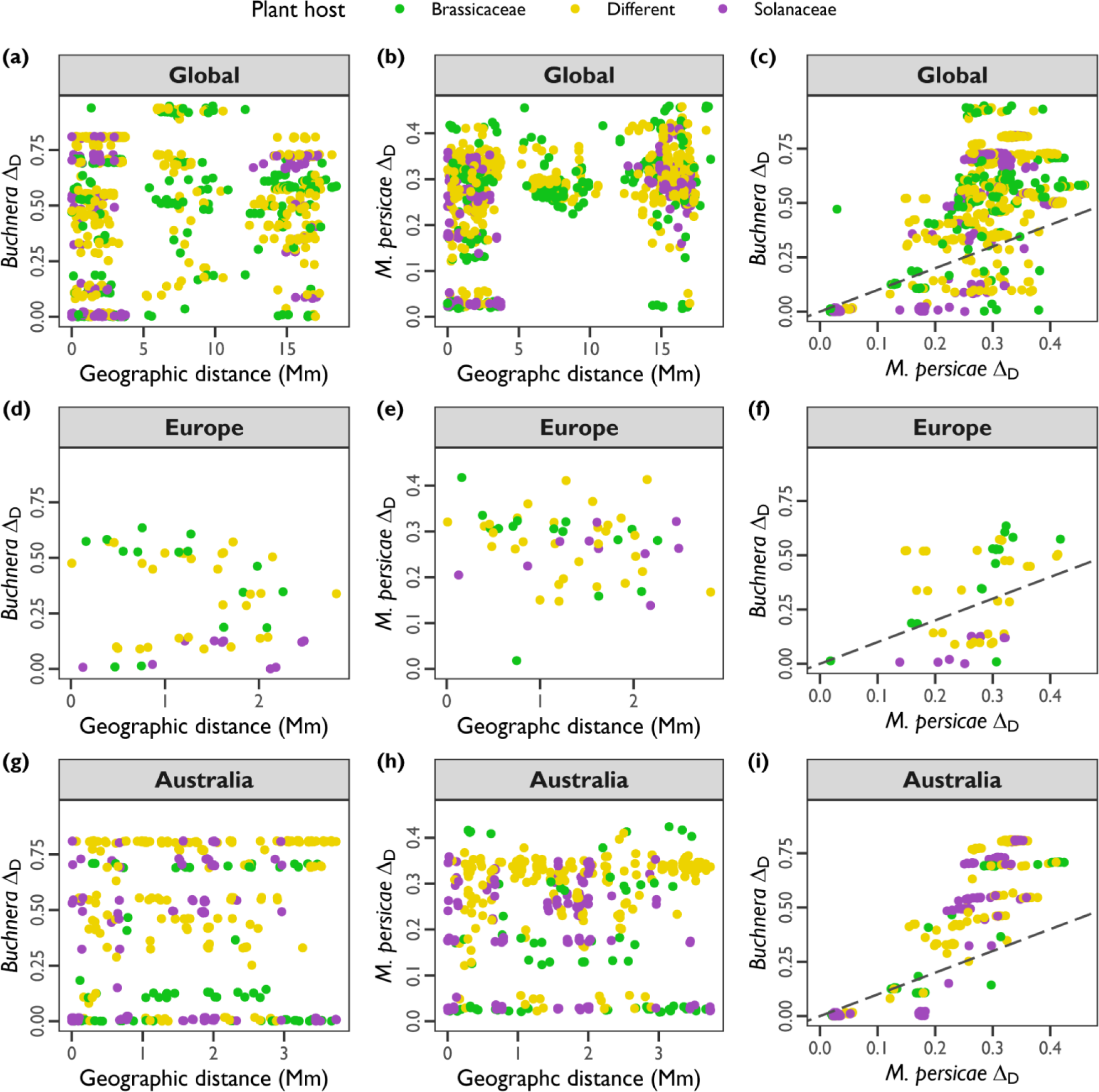
Patterns of pairwise genetic differentiation among samples. In all plots, points represent pairs of samples, colored by host plant (see legend: within Brassicaceae, within Solanaceae, or between ‘different’ plant families). Rows represent different partitions of our dataset: (a–c) global; (d–f) Europe; (g–i) Australia. Columns represent different comparisons of genetic differentiation: (a, d, g) between *Buchnera* samples (*y*-axis) as a function of geographic distance in megameters (*x*-axis); (c, e, h) between *M. persicae* samples (*y*-axis) as a function of geographic distance in megameters (*x*-axis); (c, f, i) between *Buchnera* samples (*y*-axis) as a function of genetic differentiation between *M. persicae* samples (*x*-axis), and hypothetical 1:1 relationship with dashed lines.

We did find strong support for covariation between pairs of *Buchnera* and their *M. persicae* hosts (Figures 3c, f, and i). Pearson’s correlations for pairwise Δ_D_ were *r* = 0.71, 0.37, and 0.91 in the global, European, and Australian datasets, respectively. This indicates that as aphid hosts differentiate, their *Buchnera* track them with the same trajectory of differentiation, although we found pairwise differentiation tended to be greater among *Buchnera* relative to *M. persicae* hosts. This pattern is also evidenced in the co-phylogenetic tree, where there is a general congruence of topology between *Buchnera* and their aphid host trees but the topologies are not perfectly aligned (Figure 4).

**Figure 4.**
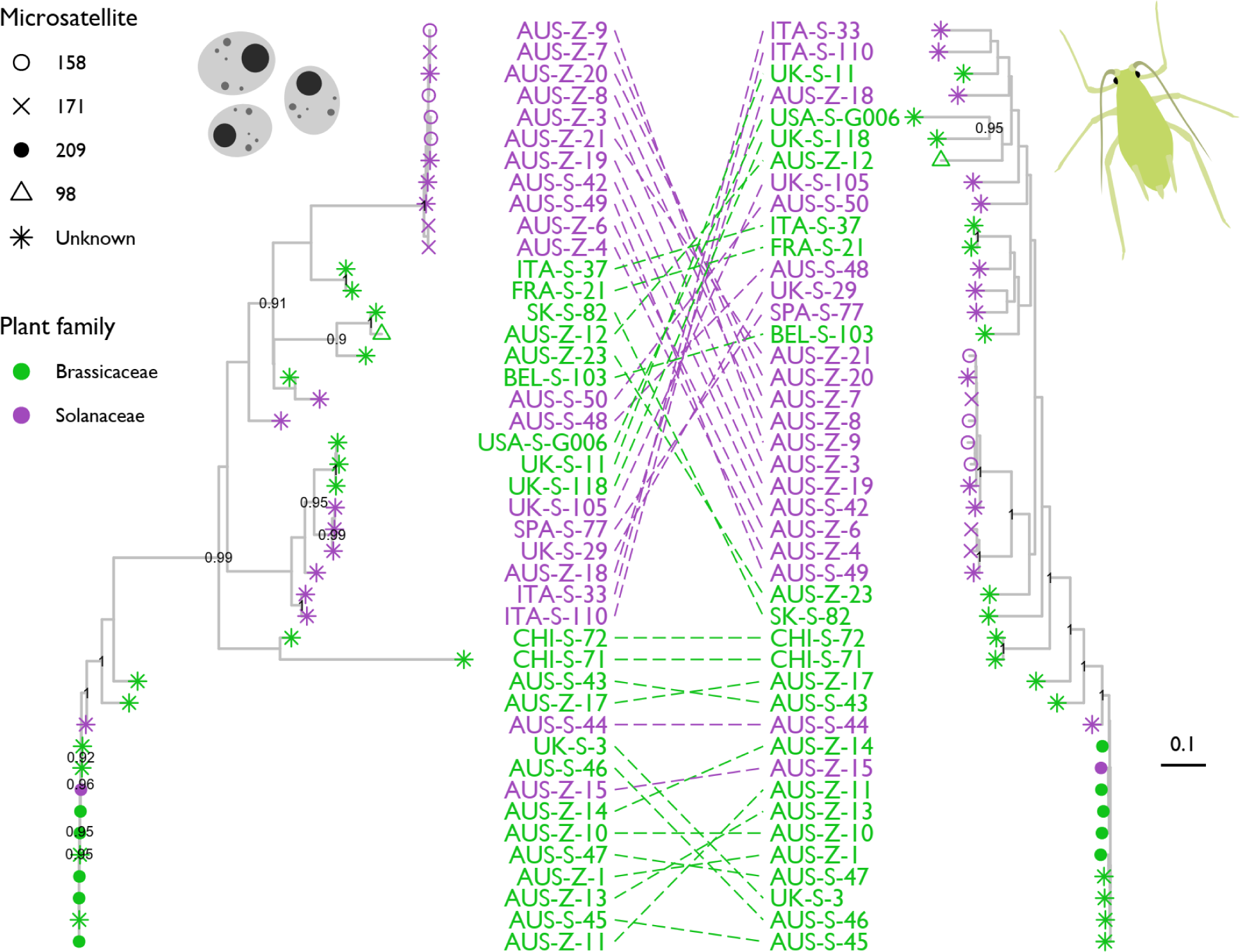
Aligned phylogenetic trees for *Buchnera* and aphid host genomes in *Myzus persicae*. The *Buchnera* tree is on the left, and the aphid host tree is on the right. Points on tree tips and sample labels are coloured by host plant; where known, post-hoc microsatellite clonal classification is depicted by point shapes (see legend). Node support is indicated as the proportion of 100 bootstrap replicates, with values >0.9 presented. The scale bar indicates the relative distance.

We did not find support for structuring of clonal strains by host plant. Residual variation in pairwise Δ_D_ was not significantly different among plant hosts at the global scale, within Europe, or within Australia (Figure 5). The exception was in the European dataset, where *Buchnera* from Solanaceae exhibited lower residual Δ_D_ relative to within Brassicaecae or different plant family comparisons, but this is likely driven by spurious effects due to small sample size (n = 10 pairs). Notably, the residual variation in Δ_D_ values was consistently distributed around 0 for all host plants, suggesting that there was no additional information in pairwise Δ_D_ values in a focal organism that could be explained after controlling for pairwise Δ_D_ values in its partner organism and geographic distance.

**Figure 5.**
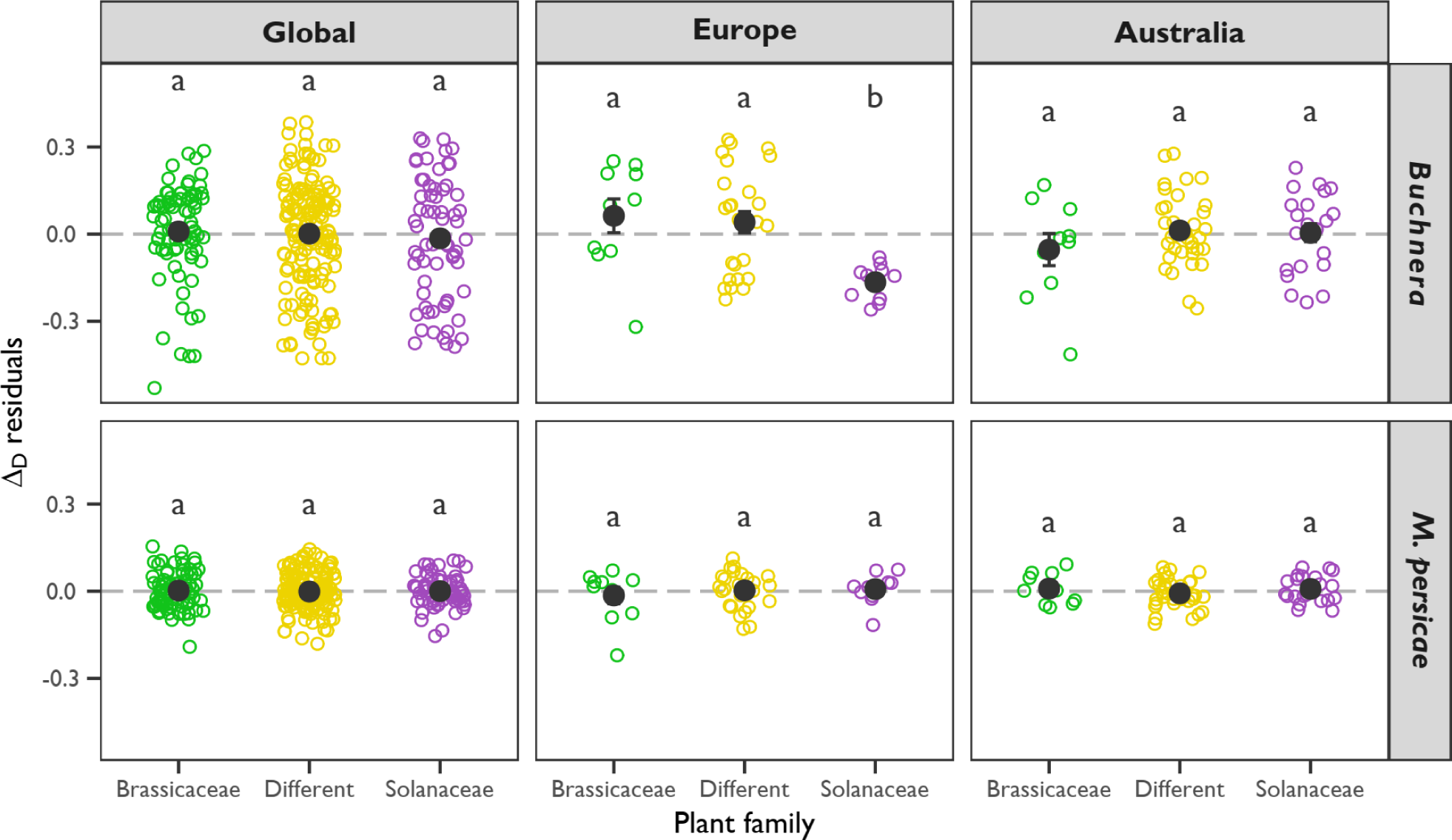
Genetic differentiation between samples of *Buchnera* or *M. persicae* from the same host plant family (within Brassicaceae or Solanaceae) or ‘different’ host plant (Brassicaceae vs. Solanaceae). Plots depict residual variation in pairwise genetic differentiation (*y*-axis) as a function of host plant (*x*-axis). Open points represent pairs of clonal samples, and large filled points represent the mean with standard error bars. Lowercase letters denote a significant difference within each group. The dashed grey line delimits a mean of zero. Each panel represents a different combination of organism (*Buchnera* or *M. persicae*, in rows) and dataset partition (global, European, or Australian, in columns).

### Mutations in *Buchnera* protein-coding regions

We recovered 1,208 biallelic variants in protein-coding regions in the *Buchnera* genome. Of these variants, 1,108 were SNPs, and 100 were complex variants, which included small microhaplotypes (multiple adjacent SNPs) or indels. Variants were annotated as 499 non-synonymous mutations and 450 synonymous mutations from coding regions, and 259 mutations from non-coding regions. Within aphid clones, we observed very strong positive correlations between the SFS of non-synonymous and synonymous *Buchnera* mutations (Supplementary Figure S2). Pearson’s correlation, *r*, had a mean of 0.98 and a range of 0.84 to 1.00 across all samples, indicating the relative allele frequency distributions of both mutation classes are very similar. However, there was a tendency for a slightly higher number of synonymous mutations relative to non-synonymous mutations (Supplementary Figure S3). Across all samples, the mean number of non-synonymous to synonymous mutations was 1.91:1, with a range of 0.63:1 and 1.24:1. There was a clear structuring of the number of mutations, and the non-synonymous and synonymous SFS, across the *Buchnera* genetic distance tree (Figure 6).

**Figure 6.**
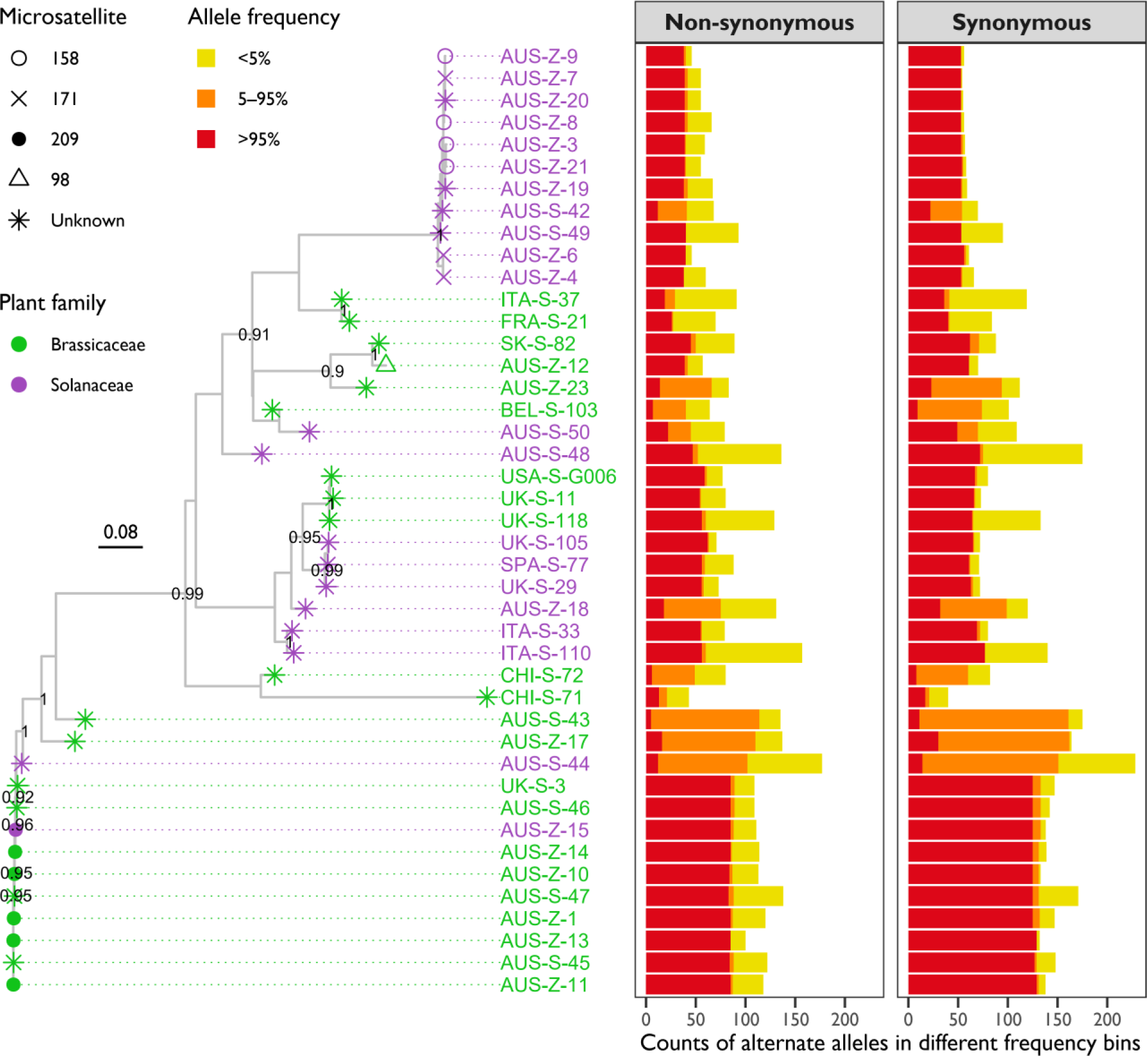
Distribution of non-synonymous and synonymous mutations along the *Buchnera* phylogenetic tree. Points on the trees tips and sample labels are colored by host plant; where known, a post-hoc microsatellite clonal classification is depicted by point shapes (see legend). Node support is indicated as the proportion of 100 bootstrap replicates, with values >0.9 presented. The scale bar indicates the relative distance. Panels depict the number of mutations (*x*-axis) per sample (*y*-axis) for non-synonymous (left) and synonymous (right) sites. Bars are portioned and colored by relative counts of different alternate allele frequency bins (<5%, 5–95%, and >95%, see legend). Classification of non-synonymous versus synonymous mutations is relative to the reference genome, GenBank Accession CP002697.1.

We tested for enrichment or diminishment of non-synonymous mutations in 68 different functional pathways across *Buchnera* samples. Following a Bonferroni correction, and significance at a threshold of *ɑ* = 0.05/68 = 0.0007, only one pathway was significant: ‘homologous recombination’, with an odds ratio of 3.06 (*p* = 0.0007), indicating an enrichment of non-synonymous mutations. If we considered significance at an uncorrected threshold of *ɑ* = 0.05, two more pathways were significant: ‘D-amino acid metabolism’, odds ratio of 0.13 (*p* =0.04), a diminishment of non-synonymous mutations; and ‘Glutathione metabolism’, odds ratio of 7.5 (*p* = 0.03), an enrichment of non-synonymous mutations.

## DISCUSSION

Macroevolutionary studies have clearly shown that *Buchnera* and their aphid hosts exhibit significant covariance in their evolutionary histories (Chen et al., 2017; Chong et al., 2019; Jousselin et al., 2009; Nováková et al., 2013). But these studies among species shed little light on the eco-evolutionary processes within species, which require a focus on microevolutionary scales. This present study bridges an important knowledge gap by examining the genomic variation and covariation in 43 clonal lineages of *M. persicae*. Our findings indicate that multiple genetically different strains of *Buchnera* can be found in a single clonal strain of *M. persicae*. This parallels very recent genomic work that has also identified coexisting *Buchnera* strains in four different clonal lineages of Russian wheat aphid, *Diuraphis noxia* (Burger, Nicolis, & Botha, 2023). Modern whole-genome sequencing clearly provides great potential to unearth previously unappreciated within and among clonal lineage variation required for asexual organisms to evolve (Loxdale & Lushai, 2003), but this has yet to be fully realized. Indeed, the limited plasticity of *Buchnera*’s genetic regulatory capacity (Moran et al., 2003; Neiers, Saliou, Briand, & Robichon, 2021; Wilcox, Dunbar, Wolfinger, & Moran, 2003) suggests that physiological effects of *Buchnera* on their aphid host is likely via genetically encoded variants, as demonstrated in the model pea aphid, *Acyrthosiphum pisum* (Chung et al., 2020; Dunbar et al., 2007; Perreau et al., 2021; Vogel & Moran, 2011). The density of *Buchnera* has largely been the focus of within-species investigations due to the importance of these endosymbionts in nutrient provisioning (Whittle, Barreaux, Bonsall, Ponton, & English, 2021), but the eco-evolutionary consequences of genetic polymorphisms within aphid hosts is virtually unknown outside the model *A. pisum*. Our results from *M. persicae* suggest that most *Buchnera* polymorphisms may not be adaptive and that microevolutionary trajectories of *Buchnera* may simply ‘drift’ with that of their aphid hosts.

We found no evidence that genetic variation in *Buchnera*, or their *M. persicae* host, was structured relative to the host plant families Brassicaceae and Solanaceae (excluding tobacco which was not considered here). The *Buchnera* genome from *M. persicae* has retained genes that may be important in the polyphagous ecology of this species (Jiang et al., 2013), yet few studies have directly examined the role of *Buchnera* in plant host adaptation in this aphid species. In the tobacco specialized subspecies, *M. p. nicotianae*, plant host adaptation appears to be completely under control of the aphid host (Jiang et al., 2013; Singh et al., 2020). Genetic variation in *Buchnera* might have little impact on the utilization of Brassicaceae and (non-tobacco) Solanaceae, at least as measured in this study.

Our more intensive sampling of invasive Australian clones recovered two genetic groups of closely related aphid clones associated with either Solanaceae or Brassicaceae, a pattern that may reflect bottlenecks in *M. persicae* following invasion into Australia (Yang et al., 2023) or perhaps selective sweeps of asexually reproducing clones that carry insecticide resistance alleles (de Little et al., 2017; Umina et al., 2014). These findings are analogous to extremely limited clonal diversity of *Sitobion avenae*, wheat aphid, in the UK, where a single asexual lineage carrying resistance to pyrethroid insecticides has risen to dominance (Morales-Hojas, Chen, Sun, Alvira, & Xiaoling, 2020). In contrast, where aphids maintain their alternating asexual–sexual life cycle, higher diversity is observed, for example, *Sitobion miscanthi* and *Schlechtendalia chinensis* in China (Morales-Hojas, Chen, et al., 2020; Sun et al., 2022; Zhang et al., 2018), or *Rhopalosiphum padi* in the UK (Morales-Hojas, Gonzalez-Uriarte, et al., 2020). Our genomic data also highlights that previous microsatellite characterizations of *M. persicae* clones in Australia (de Little et al., 2017; Umina et al., 2014) may not capture sufficient variation to reflect levels of clonal divergence. After retrospectively collecting microsatellite data to some of our sequenced clones, we found that clonal types 158 and 171 were indistinguishable from each other and did not separate into discrete monophyletic groups. Moving forward, a curated SNP genotyping panel is likely to have considerable utility in future surveys of *M. persicae* diversity and evolution in Australia.

Lack of genetic−geographic distance associations between pairs of clones highlights the dispersive and invasive nature of *M. persicae* that has spread clones across the globe. In this study, we were specifically interested in whether pairs of clones from disparate locations tend to be more genetically dissimilar than clonal samples at more proximate locations. This contrasted the goals of other genetic studies using microsatellites (Margaritopoulos et al., 2009) or genomic data (Singh et al., 2021) focusing on the beta diversity of clonal lineages among different geographic locations. We show that genetic differences between pairs of clonal *M. persicae* hosts or their *Buchnera* does not scale with geographic distance, irrespective of whether we considered the global dataset or partitions of the European or Australian clones. Indeed, we found samples in Australia that were genetically very similar to those found in Europe, suggesting few post-invasion genetic changes have occurred in *M. persicae* hosts or their *Buchnera*.

Our analysis of mutations in protein-coding regions of the *Buchnera* genome are generally consistent with genomic decay (Silva, Latorre, & Moya, 2001; Wernegreen & Moran, 1999). We suspect that neutral processes predominate the microevolutionary changes in *Buchnera* from *M. persicae*, based on our results indicating an approximate 1:1 ratio of non-synonymous to synonymous mutations (mean of 0.91:1) and similar SFS between these mutation classes. Although the exact number of mutations identified depends on the sequence divergence of a focal *Buchnera* genome from the reference genome sequence, relative patterns of mutation classes tended to be consistent across *M. persicae* clones. Most non-synonymous mutations are expected to be deleterious and can be eliminated by efficient selection within and among clones (Rispe & Moran, 2000; Wernegreen & Moran, 1999). This should bring down the ratio of non-synonymous to synonymous mutations and keep deleterious mutations at low frequencies. However, most (non-synonymous or synonymous) mutations in protein-coding regions in our *Buchnera* analysis were near or at fixation within clonal populations. These results are more consistent with a high drift scenario that drives mutations (irrespective of functional effect) to high frequency, which might be expected if the effective population size of *Buchnera* is small, selection is weak, and mutation rates are high (Bourguignon et al., 2020; Wernegreen, 2015, 2017).

Non-synonymous mutations were generally found at equivalent abundances across different functional pathways. After false positive correction, only the homologous recombination pathway was significant, and enriched for non-synonymous mutations. Homologous recombination is part of the DNA repair process, which has undergone considerable degradation in the *Buchnera* genome as it transitioned to an obligate endosymbiont (Shigenobu et al., 2000). This observation does not align with our expectations about enrichment in core cellular, core mutualism or ecological relevant pathways, nor with the expectation of neutral evolution. Without false positive correction, we also found that non-synonymous mutations were detected as being (a) diminished in D-amino acid metabolism genes, and (b) enriched in glutathione metabolism genes. Diminished non-synonymous mutations in D-amino acid metabolism may align with our core cellular pathway hypothesis. D-amino acids are used to synthesize peptidoglycan and build the cell wall. *Buchnera* from the aphid tribe Macrosiphini (which includes *M. persicae*) contribute to building their own cell wall (Smith, Li, Perreau, & Moran, 2022). Enriched non-synonymous mutations in glutathione metabolism might align with our hypothesis of ecologically relevant pathways. Prior work has suggested that glutathione metabolism in *Buchnera* may play a role in acquiring inorganic sulfur from plant phloem (Douglas, 1988; Jiang et al., 2013). Glutathione as a substrate for glutathione-S-transferases has also been reported as important for plant xenobiotic metabolism in *M. persicae* (Francis, Vanhaelen, & Haubruge, 2005), and perhaps for resistance to synthetic insecticides (Bass & Field, 2011). This suggests a possible adaptive role of glutathione metabolism in *Buchnera* supporting polyphagy and (or) insecticide detoxification in its *M. persicae* host. However, such conjectures remain speculative and neutral processes seem to predominate the genome dynamics of *Buchnera* sampled in this study from *M. persicae*.

In conclusion, our study provides a novel perspective on *Buchnera*–aphid host coevolution in *M. persicae* by focusing at the microevolutionary scale, both within and among different clones of *M. persicae*. Although our study suggests that *Buchnera* may simply ‘drift’ with their aphid host, we cannot rule out weak selection on some protein-coding changes. Whilst we largely focused on variation across clonal strains, it is still possible that there are lineage-specific variants in the *Buchnera* genome involved in coadaptation (as mutualists, antagonists, or both) (Bennett & Moran, 2015). Future experimental manipulations of *Buchnera* in different aphid host genetic backgrounds (Perreau et al., 2021) provides interesting prospects to test the eco-evolutionary consequences of genetic interactions in the *Buchnera*–aphid host symbiosis. An important extension of this work will also be to quantify variation of *Buchnera* strains within and among single aphids from the same clonal strain.

## ACKNOWLEDGEMENTS

Funding for this work came from the Grains Research Development Corporation (Australia) as part of the ‘Australian Grains Pest Innovation Program’, and from the University of Melbourne. This research was supported by The University of Melbourne’s Research Computing Services and the Petascale Campus Initiative. We thank Cesar Australia for assisting in aphid sample collections and Alex Gill for technical assistance. This manuscript was improved from the helpful comments by the Associate Editor and two anonymous referees.

## DATA AVAILABILITY

All sequencing data generated in this study has been deposited into GenBank, BioProject PRJNA987046, Sequencing Read Archive (SRA) SRR25064204 to SRR25064225. SRA Accessions for all samples used in this study are presented in Table S1, including those sourced from Singh et al. (2021). All scripts and data required to recreate these analyses are stored in a Dryad repository, doi.org/10.5061/dryad.gf1vhhmvz (Thia, 2023).

## AUTHOR CONTRIBUTIONS

JAT, KR, AAH and QY designed the research. PAU organized the sourcing of aphid material. QY organized the molecular work and sequencing. JAT led the analyses, with support from AAH, DWZ and KR. JAT wrote the original manuscript. All authors contributed to revisions of the final manuscript.

## SUPPLEMENTARY INFORMATION

**Figure S1.**
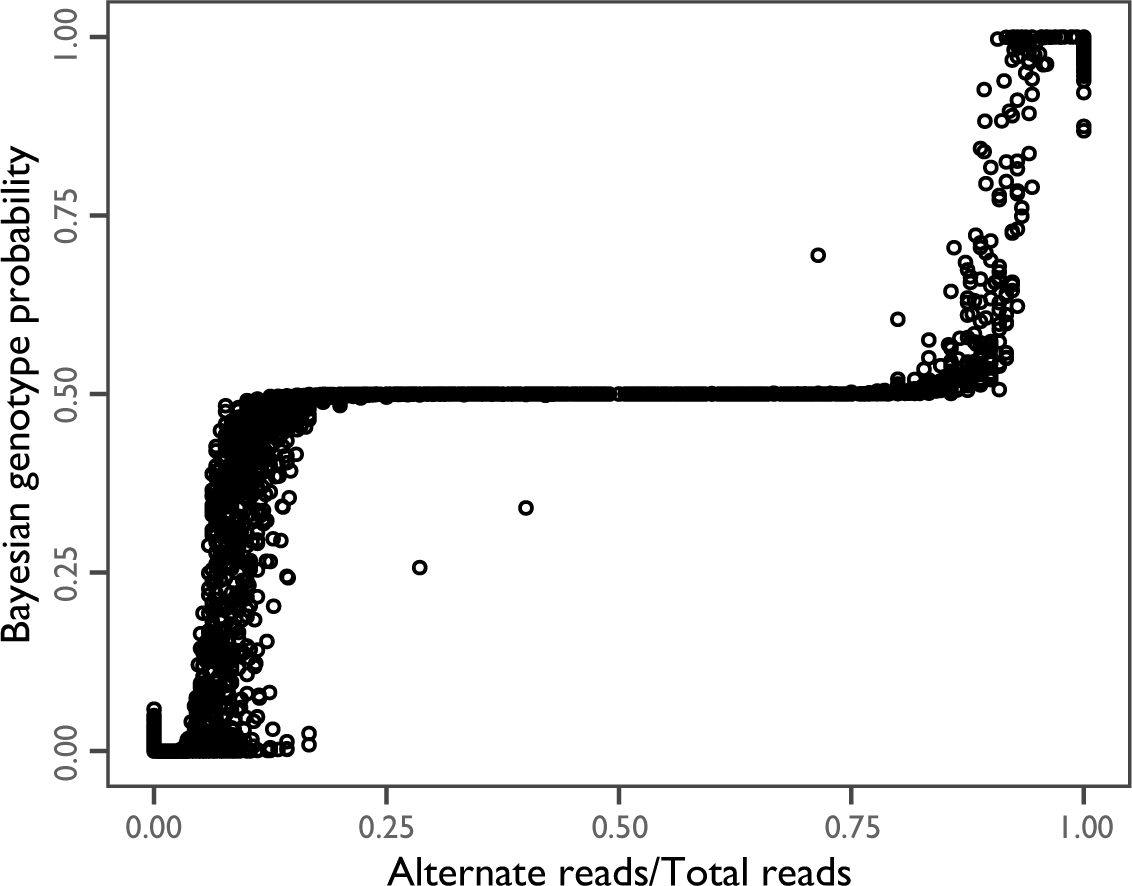
Bayesian genotype probability (*y*-axis) in clonal aphid host samples as a function of alternate read proportions (*x*-axis) as the number of alternate reads over total read depth. A genotype probability of 1 represents a homozygote for the alternate allele, a value of 0.5 represents a heterozygote, and a value of 0 represents a homozygote for the reference allele.

**Figure S2.**
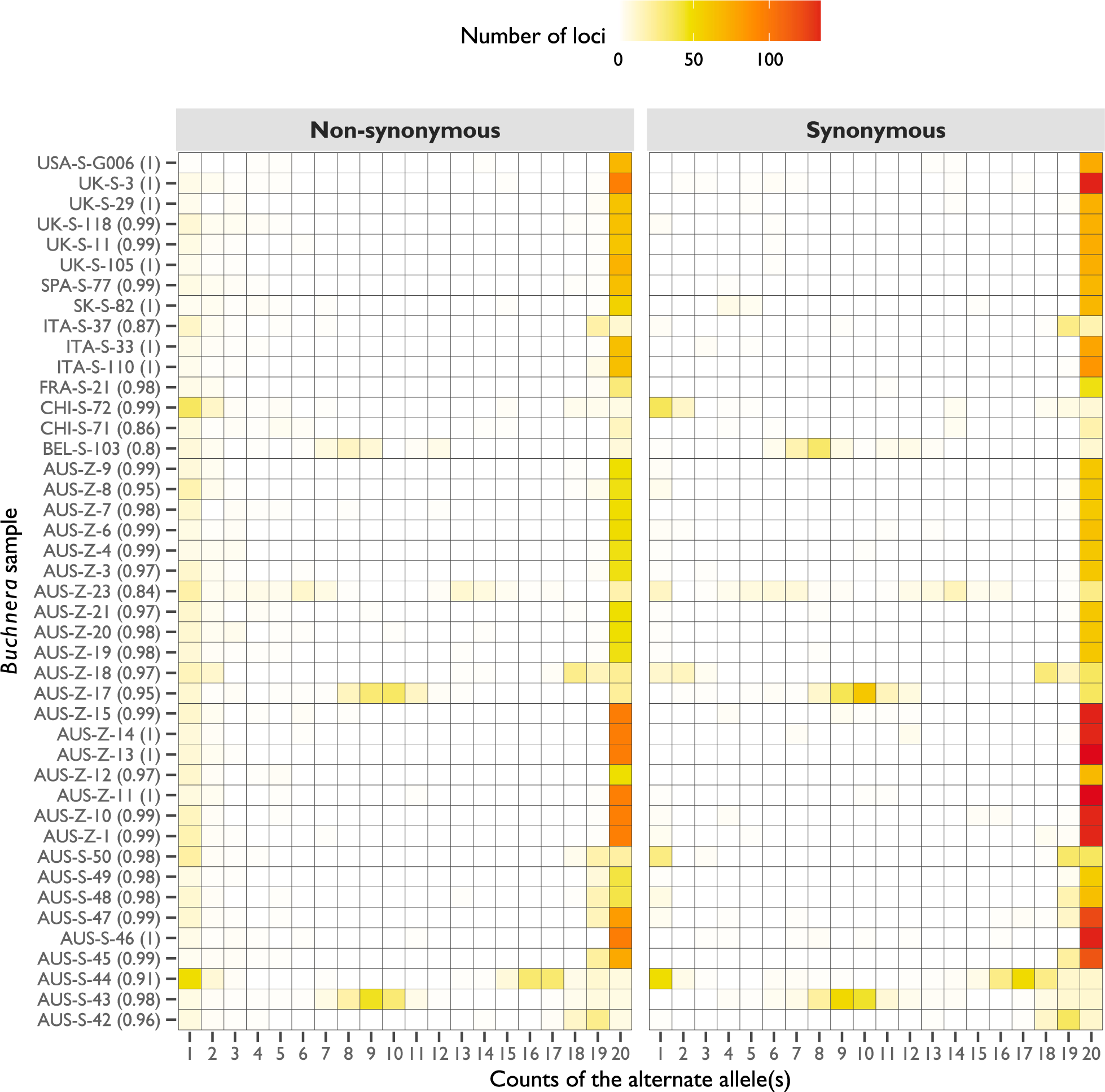
Site frequency spectra (SFS) for non-synonymous and synonymous mutations in the *Buchnera* genome. Counts of the alternate allele (*x*-axis) have been downward projected to bins of 1 to 20 for each *Buchnera* sample (*y*-axis). Values in parentheses next to each sample name indicate the Pearson correlation coefficient between non-synonymous and synonymous mutations. Cells are coloured based on the number of loci for each sample and allele count bin (see legend).

**Figure S3.**
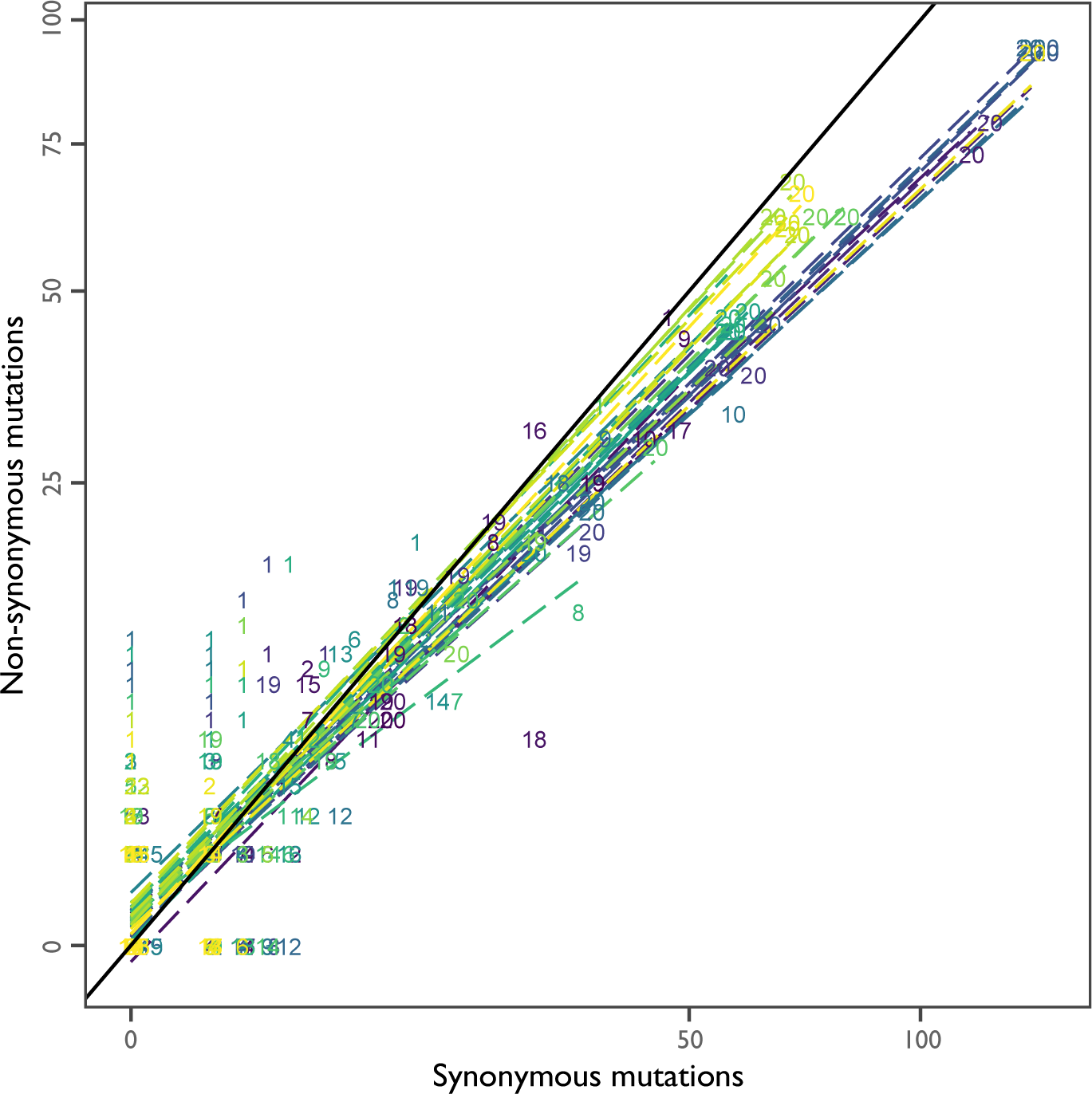
Correlations between the site frequency spectra (SFS) of non-synonymous and synonymous mutations in the *Buchnera* genome. The number of non-synonymous mutations (*y*-axis) is presented as a function of the number of synonymous mutations (*x*-axis). Numbers represent different bins of alternate allele counts (downward projected to fit in the range of 1 to 20), with dashed lines delimiting the linear relationship between mutation classes (non-synonymous vs synonymous). Numbers and dashed lines are coloured by different *Buchnera* samples. The solid black indicates a 1:1 relationship, that is, the number of non-synonymous and non-synonymous mutations for a given allele count bin are equal.

**Table S1.**
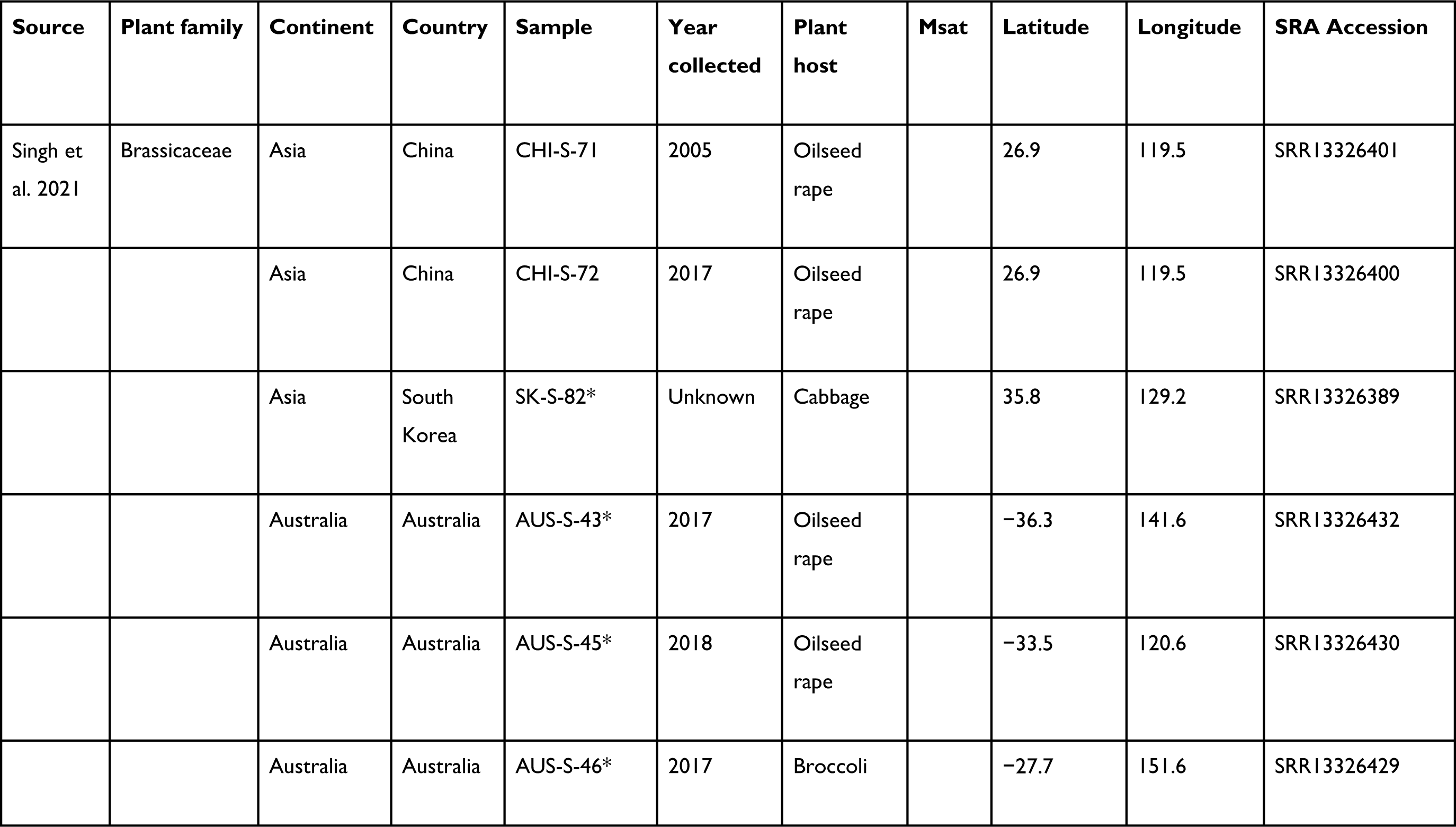
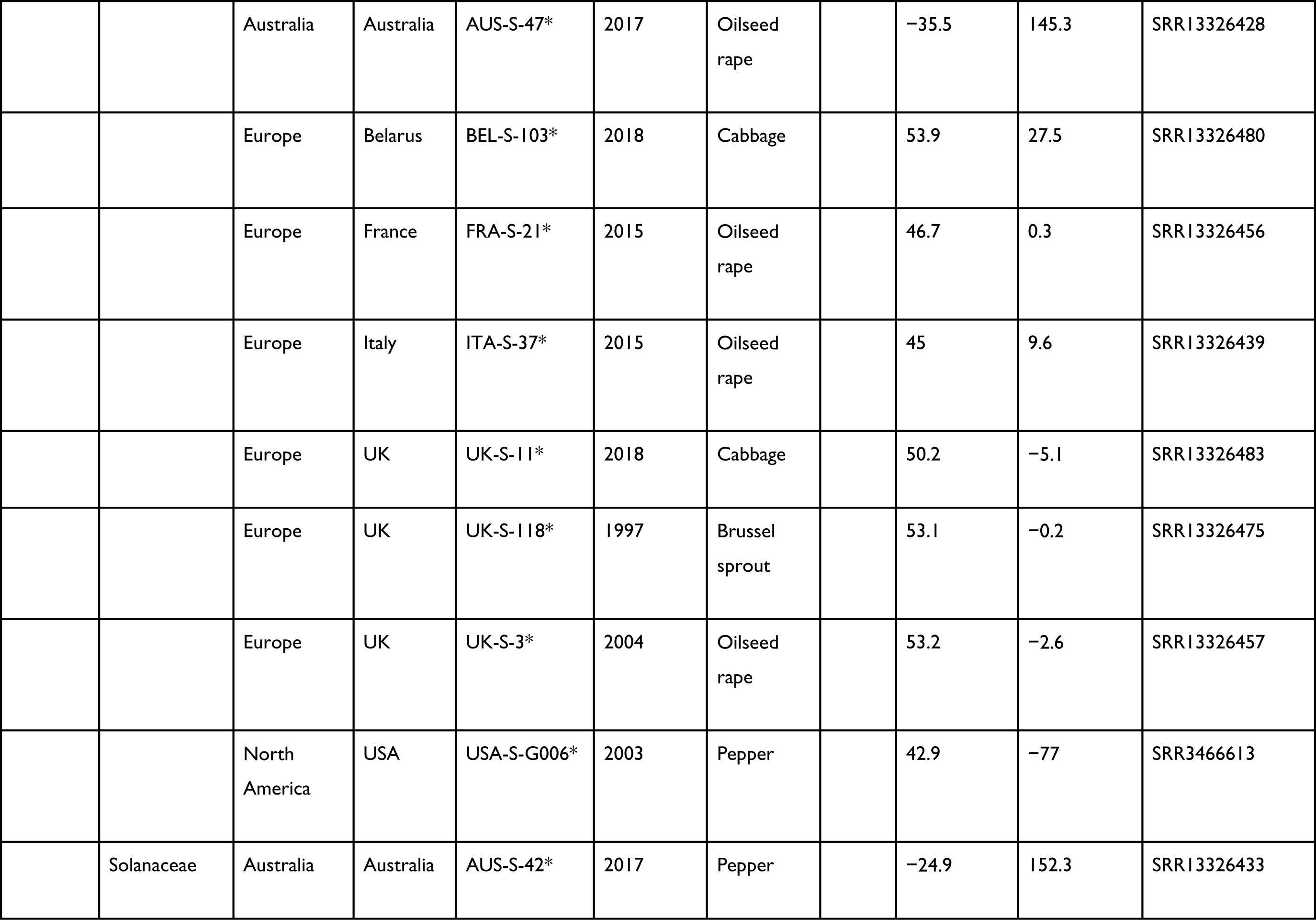
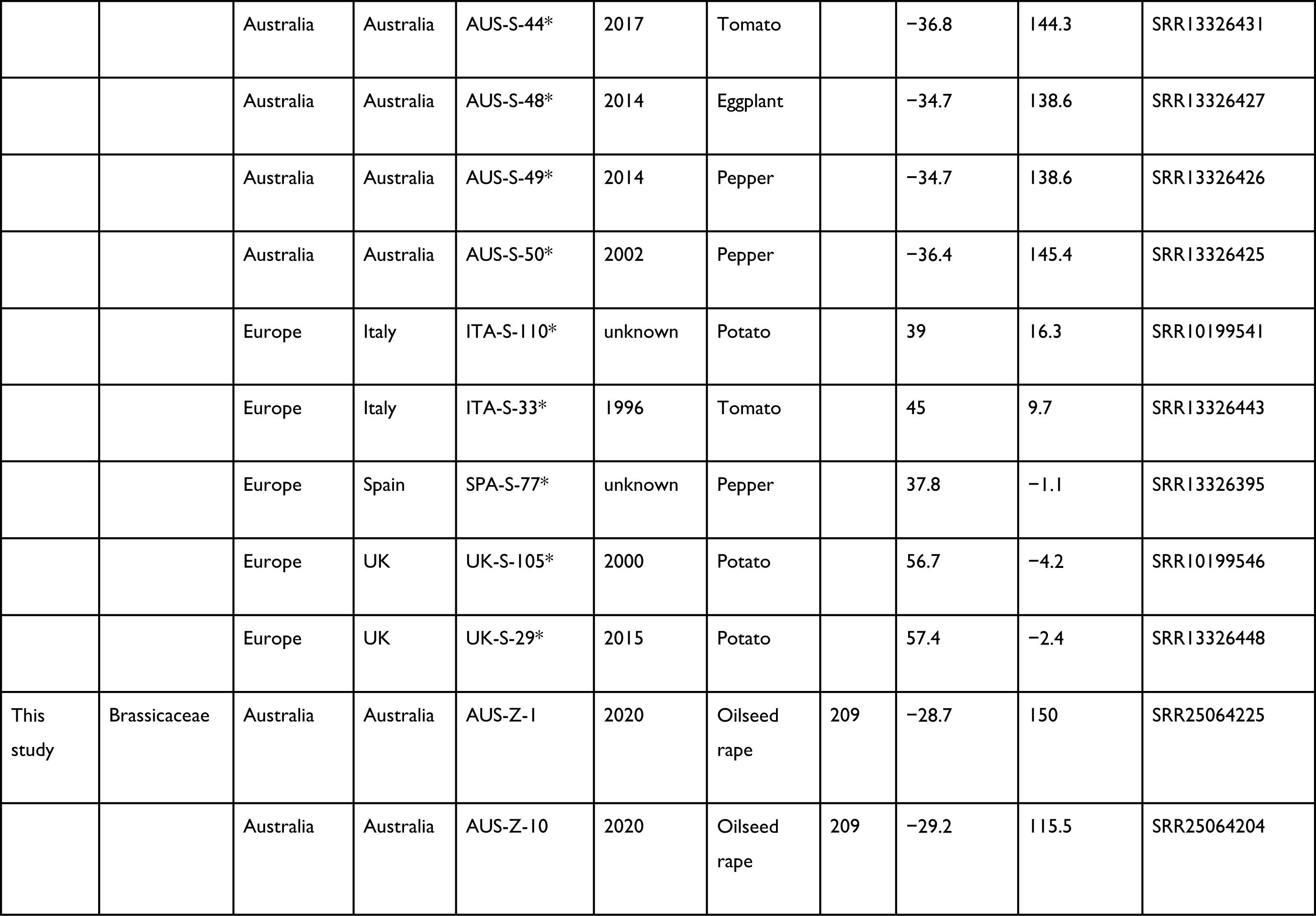
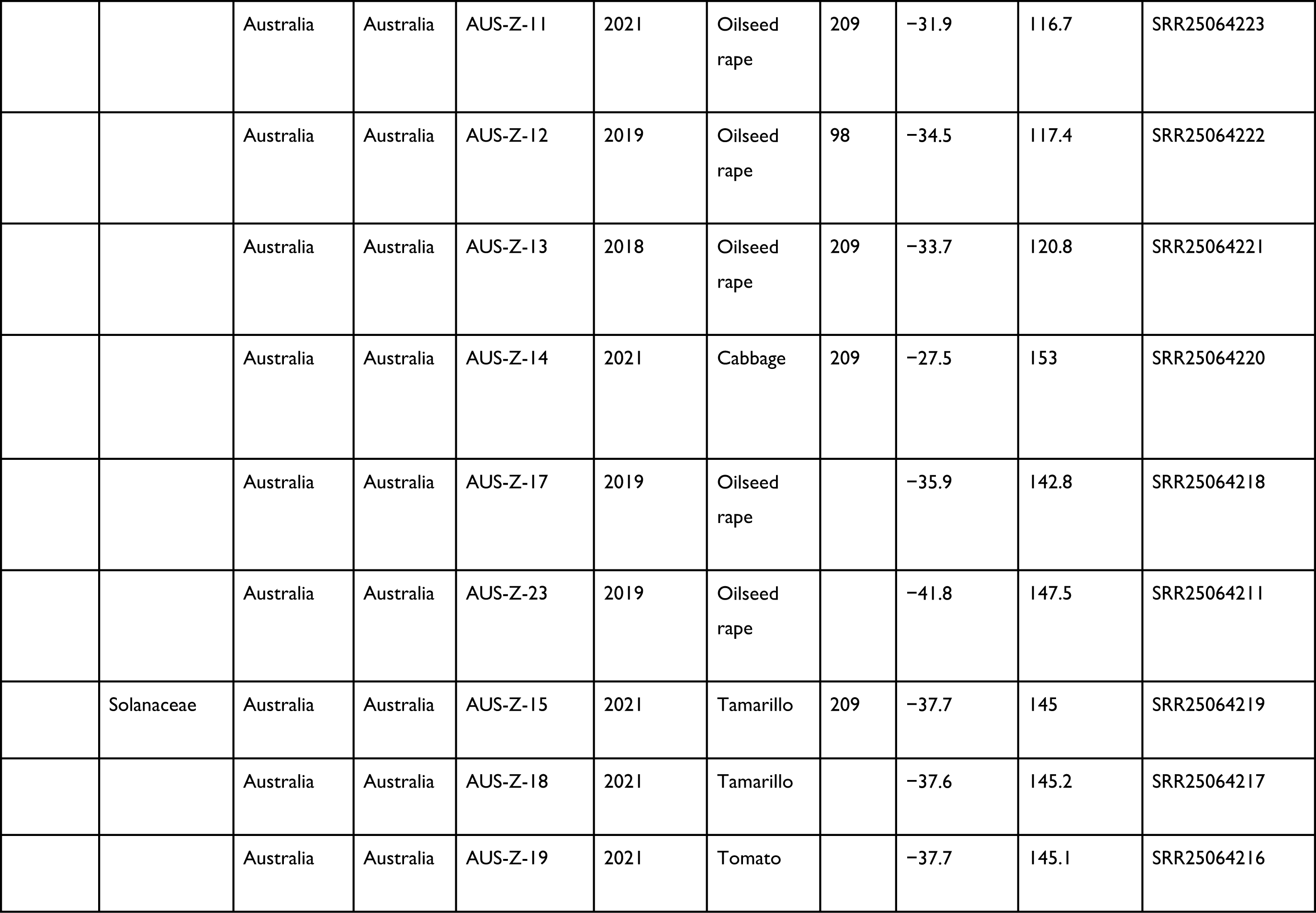
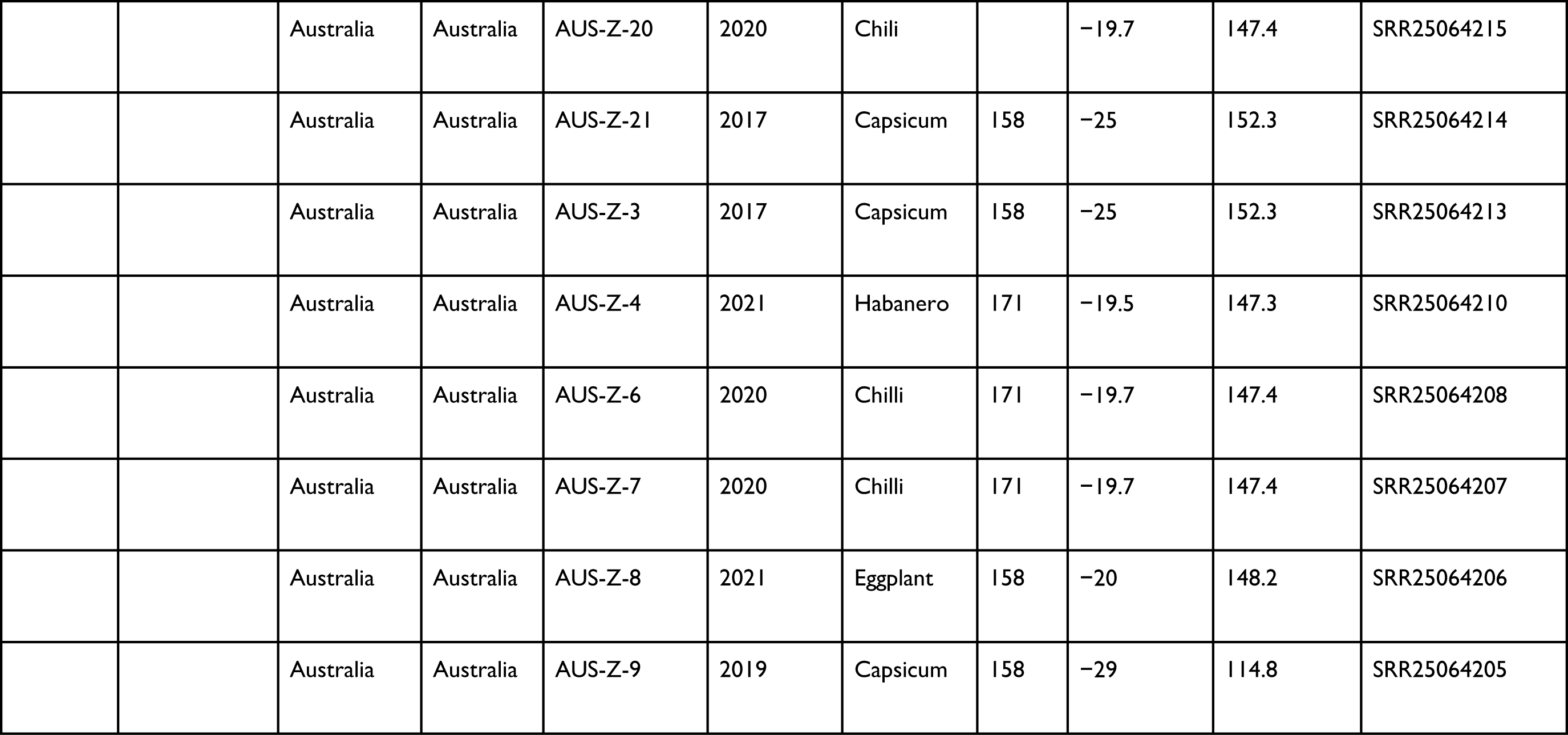
Sample information for *Myzus persicae* clones. Microsatellite assignments (Msat) are based on nomenclature from previous studies conducted on Australian *M. persicae*. Locations for some samples are approximate and based on local townships near where they were collected; these are marked with an asterisk (*).

| Source | Plant family | Continent | Country | Sample | Year collected | Plant host | Msat | Latitude | Longitude | SRA Accession |
| --- | --- | --- | --- | --- | --- | --- | --- | --- | --- | --- |
| Singh et al. 2021 | Brassicaceae | Asia | China | CHI-S-71 | 2005 | Oilseed rape |  | 26.9 | 119.5 | SRR13326401 |
|  |  | Asia | China | CHI-S-72 | 2017 | Oilseed rape |  | 26.9 | 119.5 | SRR13326400 |
|  |  | Asia | South Korea | SK-S-82* | Unknown | Cabbage |  | 35.8 | 129.2 | SRR13326389 |
|  |  | Australia | Australia | AUS-S-43* | 2017 | Oilseed rape |  | -36.3 | 141.6 | SRR13326432 |
|  |  | Australia | Australia | AUS-S-45* | 2018 | Oilseed rape |  | -33.5 | 120.6 | SRR13326430 |
|  |  | Australia | Australia | AUS-S-46* | 2017 | Broccoli |  | -27.7 | 151.6 | SRR13326429 |
|  |  | Australia | Australia | AUS-S-47* | 2017 | Oilseed rape |  | -35.5 | 145.3 | SRR13326428 |
|  |  | Europe | Belarus | BEL-S-103* | 2018 | Cabbage |  | 53.9 | 27.5 | SRR13326480 |
|  |  | Europe | France | FRA-S-21* | 2015 | Oilseed rape |  | 46.7 | 0.3 | SRR13326456 |
|  |  | Europe | Italy | ITA-S-37* | 2015 | Oilseed rape |  | 45 | 9.6 | SRR13326439 |
|  |  | Europe | UK | UK-S-11* | 2018 | Cabbage |  | 50.2 | -5.1 | SRR13326483 |
|  |  | Europe | UK | UK-S-118* | 1997 | Brussel sprout |  | 53.1 | -0.2 | SRR13326475 |
|  |  | Europe | UK | UK-S-3* | 2004 | Oilseed rape |  | 53.2 | -2.6 | SRR13326457 |
|  |  | North America | USA | USA-S-G006* | 2003 | Pepper |  | 42.9 | -77 | SRR3466613 |
|  | Solanaceae | Australia | Australia | AUS-S-42* | 2017 | Pepper |  | -24.9 | 152.3 | SRR13326433 |
|  |  | Australia | Australia | AUS-S-44* | 2017 | Tomato |  | -36.8 | 144.3 | SRR13326431 |
|  |  | Australia | Australia | AUS-S-48* | 2014 | Eggplant |  | -34.7 | 138.6 | SRR13326427 |
|  |  | Australia | Australia | AUS-S-49* | 2014 | Pepper |  | -34.7 | 138.6 | SRR13326426 |
|  |  | Australia | Australia | AUS-S-50* | 2002 | Pepper |  | -36.4 | 145.4 | SRR13326425 |
|  |  | Europe | Italy | ITA-S-110* | unknown | Potato |  | 39 | 16.3 | SRR10199541 |
|  |  | Europe | Italy | ITA-S-33* | 1996 | Tomato |  | 45 | 9.7 | SRR13326443 |
|  |  | Europe | Spain | SPA-S-77* | unknown | Pepper |  | 37.8 | -1.1 | SRR13326395 |
|  |  | Europe | UK | UK-S-105* | 2000 | Potato |  | 56.7 | -4.2 | SRR10199546 |
|  |  | Europe | UK | UK-S-29* | 2015 | Potato |  | 57.4 | -2.4 | SRR13326448 |
| This study | Brassicaceae | Australia | Australia | AUS-Z-1 | 2020 | Oilseed rape | 209 | -28.7 | 150 | SRR25064225 |
|  |  | Australia | Australia | AUS-Z-10 | 2020 | Oilseed rape | 209 | -29.2 | 115.5 | SRR25064204 |
|  |  | Australia | Australia | AUS-Z-11 | 2021 | Oilseed rape | 209 | -31.9 | 116.7 | SRR25064223 |
|  |  | Australia | Australia | AUS-Z-12 | 2019 | Oilseed rape | 98 | -34.5 | 117.4 | SRR25064222 |
|  |  | Australia | Australia | AUS-Z-13 | 2018 | Oilseed rape | 209 | -33.7 | 120.8 | SRR25064221 |
|  |  | Australia | Australia | AUS-Z-14 | 2021 | Cabbage | 209 | -27.5 | 153 | SRR25064220 |
|  |  | Australia | Australia | AUS-Z-17 | 2019 | Oilseed rape |  | -35.9 | 142.8 | SRR25064218 |
|  |  | Australia | Australia | AUS-Z-23 | 2019 | Oilseed rape |  | -41.8 | 147.5 | SRR25064211 |
|  | Solanaceae | Australia | Australia | AUS-Z-15 | 2021 | Tamarillo | 209 | -37.7 | 145 | SRR25064219 |
|  |  | Australia | Australia | AUS-Z-18 | 2021 | Tamarillo |  | -37.6 | 145.2 | SRR25064217 |
|  |  | Australia | Australia | AUS-Z-19 | 2021 | Tomato |  | -37.7 | 145.1 | SRR25064216 |
|  |  | Australia | Australia | AUS-Z-20 | 2020 | Chili |  | -19.7 | 147.4 | SRR25064215 |
|  |  | Australia | Australia | AUS-Z-21 | 2017 | Capsicum | 158 | -25 | 152.3 | SRR25064214 |
|  |  | Australia | Australia | AUS-Z-3 | 2017 | Capsicum | 158 | -25 | 152.3 | SRR25064213 |
|  |  | Australia | Australia | AUS-Z-4 | 2021 | Habanero | 171 | -19.5 | 147.3 | SRR25064210 |
|  |  | Australia | Australia | AUS-Z-6 | 2020 | Chilli | 171 | -19.7 | 147.4 | SRR25064208 |
|  |  | Australia | Australia | AUS-Z-7 | 2020 | Chilli | 171 | -19.7 | 147.4 | SRR25064207 |
|  |  | Australia | Australia | AUS-Z-8 | 2021 | Eggplant | 158 | -20 | 148.2 | SRR25064206 |
|  |  | Australia | Australia | AUS-Z-9 | 2019 | Capsicum | 158 | -29 | 114.8 | SRR25064205 |

